# Individual Differences in Training Naive Listeners to Localize Spatial Audio in Virtual Reality

**DOI:** 10.1101/2025.10.25.681945

**Authors:** Tanya Wen, Antje Ihlefeld

## Abstract

It has been widely believed that a key factor in creating realistic spatial audio in virtual reality (VR) is the head-related transfer function (HRTF), which is unique to each individual, but costly to measure for widespread use. This study investigates the effects of HRTF personalization and training on sound localization accuracy in VR. Two experiments were conducted: Experiment 1 compared naive listeners and those who underwent brief training on localization tasks using personalized versus generic HRTFs; Experiment 2 used a within-subject design to assess training effects over two sessions. Results show that accurately localizing sound can be a difficult task for many participants the first time. Training significantly improves localization accuracy, reducing errors and confusions, and enabling many initially non-sensitive listeners to perceive spatial audio effectively. Although HRTF personalization yielded a statistically significant benefit, the effect was small, primarily improving elevation perception at extreme angles. These findings suggest that generic HRTFs combined with user training may suffice for most VR applications.

## Introduction

Binaural rendering is used to reproduce realistic 3D auditory scenes. As the popularity of head-mounted virtual reality (VR) and augmented reality (AR) devices has increased, so has the desire for simulating realistic audio. Spatial audio gives the listener the sensation that sound is coming from the 3D environment, helps orient the listener in space, and also establishes immersion in space by providing congruous input with the visual surround (Potter et al., 2022; Rafaely et al., 2022). Users can feel more present and connected to their virtual experiences, whether that is an action-packed VR game, a group video call on a VR headset, or benefiting from audio-guided navigation on smart glasses.

It has been suggested that a key component of reproducing spatialized audio is the head-related transfer function (HRTF). The HRTF is a filter that describes how sound is transformed from the source to the listener’s ears, and it is unique to each individual, as one’s head, ears, and torso uniquely affect the acoustic path of the sound before it reaches the ear canal (Møller et al., 1995). To obtain an individual’s HRTF, the person would need to sit still in an anechoic chamber with in-ear microphones to record the sounds played from various directions. On the other hand, there are existing generic HRTFs available, that were obtained using a manikin or a donor HRTF, and may or may not include additional processing (Vorländer, 2004). Multiple prior studies have demonstrated that participants show improved performance in localization tasks using their personalized HRTF compared to a non-individualized one (Ben-Hur et al., 2020; Larsen et al., 2013; Middlebrooks, 1999; Møller et al., 1996; Rummukainen et al., 2021).

The use of non-individualized HRTFs often result in increased localization errors, such as front-back, elevation compression, and lack of externalization (Jenny & Reuter, 2020; Møller et al., 1996; Simon et al., 2016; Wenzel et al., 1993). However, measuring individualized HRTFs is a cumbersome and expensive process, making it impractical to scale to product.

On the other hand, the human auditory system exhibits high plasticity (Keating & King, 2015; King et al., 2001). For example, manipulating acoustic cues by altering the shape of the pinna or blocking one ear, participants initially show a decreased localization ability, but improve over the course of the next few days (Carlile et al., 2014; Hofman et al., 1998). Subsequent studies have shown that users can adapt to non-individualized HRTFs through continuous exposure as well as training (Parseihian & Katz, 2012; Poirier-Quinot & Katz, 2021; Shinn-Cunningham et al., 1998; Stitt et al., 2019). Training does not necessarily need to be extensive, even a small number of short (12-minute) training sessions on virtual sound can show a significant improvement on non-individualized HRTFs, which can be retrained across multiple days (Steadman et al., 2019). Some studies have even suggested that due to the adaptive nature of the auditory system, generic HRTFs may be enough to enable good auditory source localization in VR, such that there is no need for personalization (Berger et al., 2018; Lladó, Pollack, et al., 2024).

While training has been effective, we note that one study showed limited effectiveness of HRTF training in VR (Poirier-Quinot et al., 2024), with reasons to be speculated. Possible factors may include the frequency responses of the headphones, poor match between participants’ and the to-be-learned HRTF, and unlucky participant selection. Previous studies suggested that individual differences in learning a new HRTF vary greatly between individuals, and that a certain percentage of non-proficient learners should be expected in HRTF learning experiments (Parseihian & Katz, 2012; Poirier-Quinot et al., 2022; Stitt et al., 2019). For example, Stitt et al. (2019) observed that while some subjects were able to localize using non-individualized HRTFs to the same degree as their own after training, others showed little to no improvement even after 10 training sessions, including failure to resolve front-back confusions.

Here, we investigated the effect of HRTF personalization (pHRTF) on localization accuracy, compared to a generic HRTF from the Meta XR Audio SDK (uHRTF). In the first experiment, we tested two groups of participants on a sound localization task. One group (**Naive**), performed the task without any prior exposure or training, while the other group (**Train**) first underwent a training task with visual feedback. To examine the effect of training within the same subjects, in the second experiment (**Naive + Train**), we recruited naive participants to perform sound localization without prior training in the first session, and explored the effect of training on a second session on a different day.

### Experiment 1

In Experiment 1, we sought to experimentally capture the effect of an out-of-the-box experience when experiencing spatial audio in VR for the first time. In the first group of participants we tested, who were all naive to our experiment, localized sound in VR using a generic HRTF and their own personal HRTF. If HRTF personalization is the key to reproducing realistic audio, users should be able to perceive sound as they would in the real-world, and they should be more accurate in performance than using a generic HRTF. Given previous literature on the plasticity of the auditory system to learn new HRTFs, we also investigated the effect of training, where we provided visual feedback after each localization trial. While some studies suggested that it may take up to six weeks to learn an altered HRTF (Hofman et al., 1998), others suggest that only a few minutes may be sufficient (Berger et al., 2018; Mendonça et al., 2012). Here, we conducted the same localization test on a second group of participants, but implemented a short training block just prior to the localization test. If training was beneficial to localization performance, we expect participants who underwent training to perform better than the naive group, in both HRTF conditions.

## Methods

### Participants

A total of twenty-nine participants were included in Experiment 1. Twelve participants were included in the “**Naive**” group (ages: 27-44, mean = 35.5, SD = 7.43; 2 males, 10 females). Three additional participants were excluded as they were observed to not follow instructions (holding the controllers in their laps and clicking through the entire time), and two additional participants were excluded due to technical issues. In the “**Train**” group, seventeen different participants were included (ages: 23-64, mean = 38.29, SD = 12.25; 9 males, 8 females). Two additional participants were excluded due to self-reported hearing or neurological issues, and one additional participant was excluded due to technical difficulties. All participants that were included self-reported to have normal hearing and had their HRTF previously measured in our anechoic chamber within the last two years. Data were collected at Meta Reality Labs, and the study was carried out in accordance with ethical approval obtained from the Institutional Review Board. All participants provided informed consent for their participation in the study. The data and code have been made publicly available and can be accessed at https://github.com/facebookresearch/AuditoryLocalizationTraining.

### Experimental Design and Stimuli

Participants performed an auditory source localization test, where they were asked to identify the source of a 2-second pink “burst noise” sound (10 ms on/off ramps, 0.1-18 kHz cutoff) presented at 75 dB SPL, that was rendered using the Quest 3 headset, and played out through DT990 headphones. The equipment used for the study is shown in Figure 1. The headphones were equalized using the Brüel and Kjær HATS 5128-B mannikin and the inverse equalization filter was convolved with the audio from the Quest 3. Additionally, either the participant’s personal HRTF (i.e., pHRTF) or the Universal HRTF (i.e., uHRTF) (https://developers.meta.com/horizon/blog/improve-spatial-audio-universal-hrtf-meta-quest/) within the Meta XR Audio SDK (https://developers.meta.com/horizon/documentation/unity/meta-xr-audio-sdk-features) HRTF was convolved with the audio, to create spatialized audio. Sounds were rendered at a fixed 2 meter distance, at azimuths [−150°, −120°, −90°, −60°, −30°, 0°, 30°, 60°, 90°, 120°, 150°, 180°] and elevations [−60°, −30°, 0°, 30°, 60°], resulting in a total of 60 locations per block.

**Figure 1.**
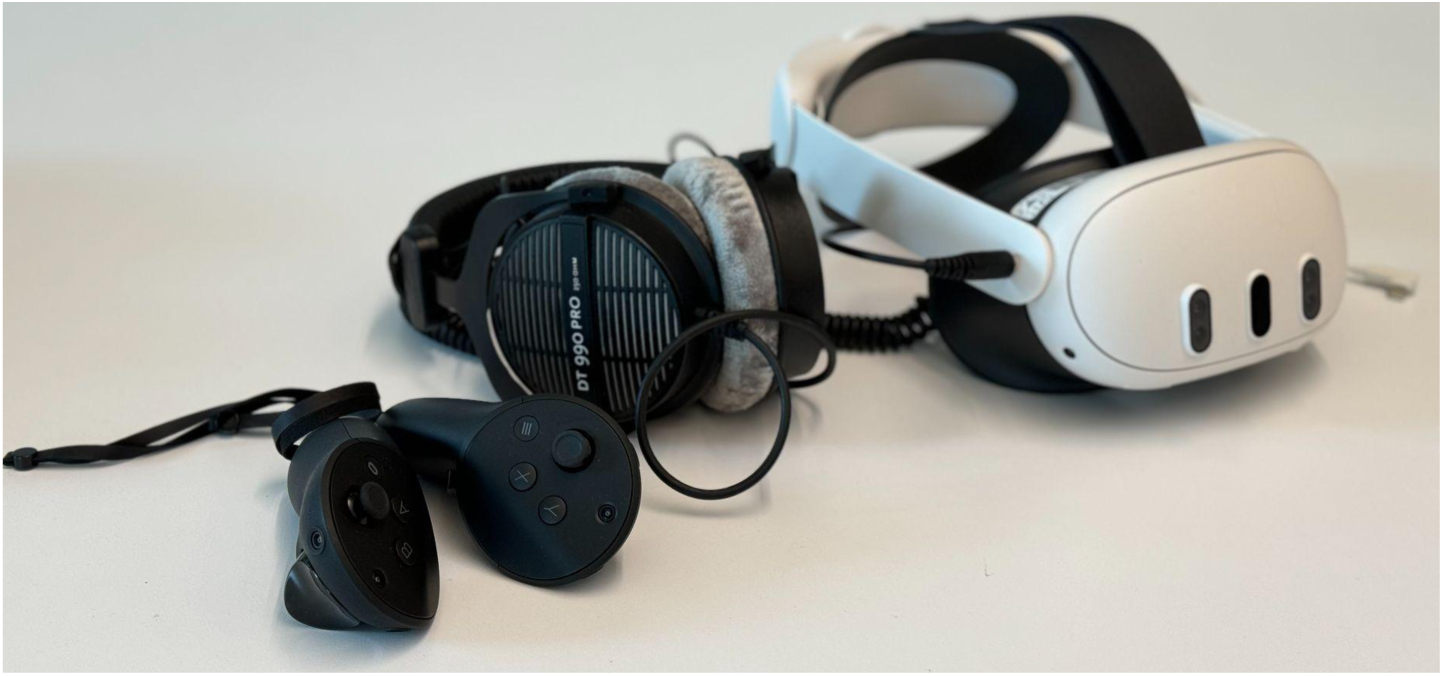
Equipment used during the sound localization test. From left to right: Quest Pro controllers, DT990 headphones, and Quest 3 headset with elite strap. During the experiment sounds were rendered though the Quest 3 headset, while participants were situated in VR and performed a localization task using the Quest Pro controllers.

To make the results of this study applicable to real-world listening in potential products, the experiment was conducted in a small everyday office room. During the main experiment, participants were situated in a gray sphere inside VR. Before the beginning of each block, participants were asked to sit facing forward in a comfortable position with their head upright. A button on the controller was pushed to confirm this as the alignment position. During the experiment, participants were given a real-time head feedback GUI, and were required to be aligned before the start of each trial. Once the sound started playing on each trial, participants were allowed to make small head movements as if they were naturally listening (no more than approximately ±10-15°). The aim of this dynamic listening design was to allow participants to use the binaural dynamic cues, while evaluating localization accuracy in the source direction that was being tested (Ben-Hur et al., 2020). Once participants had located the sound, they were allowed full range of motion, and were instructed to reach out with the left or right Quest Pro controller to indicate the sound location, by pressing the “X” or “A” button. The button position was recorded as their response location. No feedback on their localization accuracy was given to the participants in the main experiment. Participants performed a total of 4 blocks, with 2 blocks rendered with their personalized HRTF and 2 blocks rendered with the Audio SDK in a randomized order, which was not revealed to participants. Each block took approximately 7 minutes to complete.

Participants in the **“Naive”** group started the sound localization task immediately after the instructions of the experiment were given. We note that due to this being an earlier experimental session, some of the participants were only presented with azimuths from - 120° to 120° and performed a shorter session. The participants were informed that the sounds can come from any direction of the dome, and were given examples, such as “top right in front”, “lower left in the back”, and were encouraged to consider the entire response space.

Those in the **“Train”** group were given an extra block at the beginning of the study to learn to localize a 3-second pink “random click” noise that was temporally more structured as compared to the burst noise. The sound locations were jittered [−5° to 5°] around azimuths = [45°, 135°, 225°, 315°] and elevations = [−70°, −40°, −10°, 10°, 40°, 70°].

We additionally added 12 trials of random locations drawn from azimuths = [0° to 359°] and elevations = [−80° to 80°]. This resulted in a total of 60 training trials. During the training task, as soon as participants indicated the perceived location of the sound, the actual sound source location was revealed as a green dot. Participants were asked to look at the green dot and point the controller to confirm its location. Half of the participants were trained on the uHRTF and the other half were trained on their own pHRTF. The **“Train”** group then performed the same localization task as the **“Naive”** group.

### Statistical analysis

To compare the effects of personalization and training, we identified the following measures of interest: (1) localization acuity for azimuth and (2) elevation, calculated by comparing the actual azimuth elevation of the sound source with the perceived azimuth and elevation indicated by the participants; (3) arc errors, defined as the distance between the participant’s response and the virtual sound location on the sphere in units of degrees (Poirier-Quinot et al., 2022); (4) confusion errors including front/back and (5) top/down confusions, indicated by whether a sound is misclassified in hemispheric location; and (6) response time, defined as the onset of the sound to when the participant submits a response with the controller. Before fitting statistical models on response time, outliers greater than three standard deviations from the mean of each participant were removed.

Linear mixed effects model (LME) fitted the measurement of interest by maximum likelihood, using full interaction models, with Azimuth (α_y_*_1_*), and Elevation (α_y*2*_), HRTF (α_y*3*_), Group (α_y*4*_) as fixed effects. For simplicity, and comparable representation as elevation, azimuths behind the participant were folded into the front (e.g., 120° would be analyzed as 30°). Individual differences in the overall range of elevations were modeled via random effect β_y0_. The LME is described as in Equation 1. All data processing was performed using custom scripts in Matlab (Mathworks, Natick, MA).

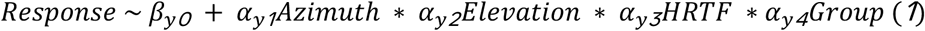

As not all participants may be sensitive to spatial audio; next, we examined the effect of HRTF personalization and training in the listeners who were deemed sensitive, using the same model above. To define sensitive listeners in an unbiased manner, we identified four quadrants in space where the participant must have indicated a response when localizing with their own HRTF. For azimuth, a response must be made in both hemispheres ranging from [−120° to 0°] and [0° to 120°]. We chose 120° as some earlier participants were not presented with further azimuths. Critically, participants must show elevation sensitivity, where responses must be made sufficiently below and above horizon, defined as between [−30° to −90°] and [30° to 90°]. This classification of sensitive and non-sensitive participants showed high agreement with human classification (Cohen’s Kappa = 0.93).

## Results

### Participant characteristics

Within the “**naive**” group, 7 out of the 12 participants (58.33%) were labeled as non-sensitive to spatial audio. In the “**train**” group, only 2 out of the 17 participants (11.76%) were classified to be non-sensitive. A Chi-squared test revealed a significant association between listener sensitivity and group (χ² = 7.13, p < 0.01).

Figure 2 shows some examples of characteristic non-sensitive listeners, as well as a typical good listener. We observed a large degree of variation in localization accuracy among participants in both groups. While some participants seem able to accurately perceive spatial audio in various directions even without training, some trained participants still remained non-sensitive. Among non-sensitive participants, we observed systematic patterns of errors, including participants exhibiting (1) lacked elevation sensitivity (sounds perceived bring mostly around horizon; N=3), (2) perceived sounds mostly around or above horizon (N=4), (3) perceived sounds largely from the back (N=3), and/or (4) discriminated sounds mostly only left versus right (N=2).

**Figure 2.**
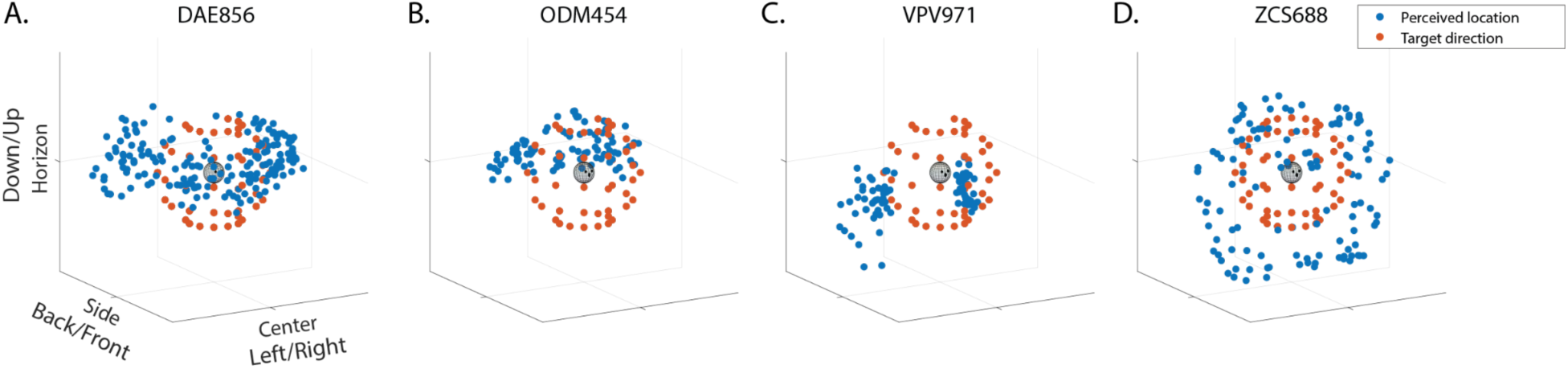
Example responses of four participants in the “naive” group while localizing using their own pHRTF. The red dots indicate the direction of the source source (scaled at 20% of the original distance for visualization). The blue dots indicate participants’ responses, with each dot representing a response for each trial. (A) Participant DAE856 showed a lack of elevation range in their responses; (B) ODM454 localized sounds with a strong bias towards the back and perceived them largely around or above horizon; (C) VPV971 showed a strong lateral bias, discriminating mainly left versus right in the horizontal plane; (D) ZCS688 is a representative example of a sensitive listener, perceiving sounds from all directions.

### Azimuth Localization

Figure 3 illustrates the localization of performance of azimuth, collapsed across elevations, in the “**naive”** and “**train”** groups when sounds were rendered with the uHRTF or their pHRTF, respectively. Individual participants are depicted as hollow circles (sensitive listeners) and asterisks (non-sensitive listeners), with each participant staggered from left to right and indexed by size. Group averages of the sensitive listeners are illustrated with the solid gray line.

**Figure 3.**
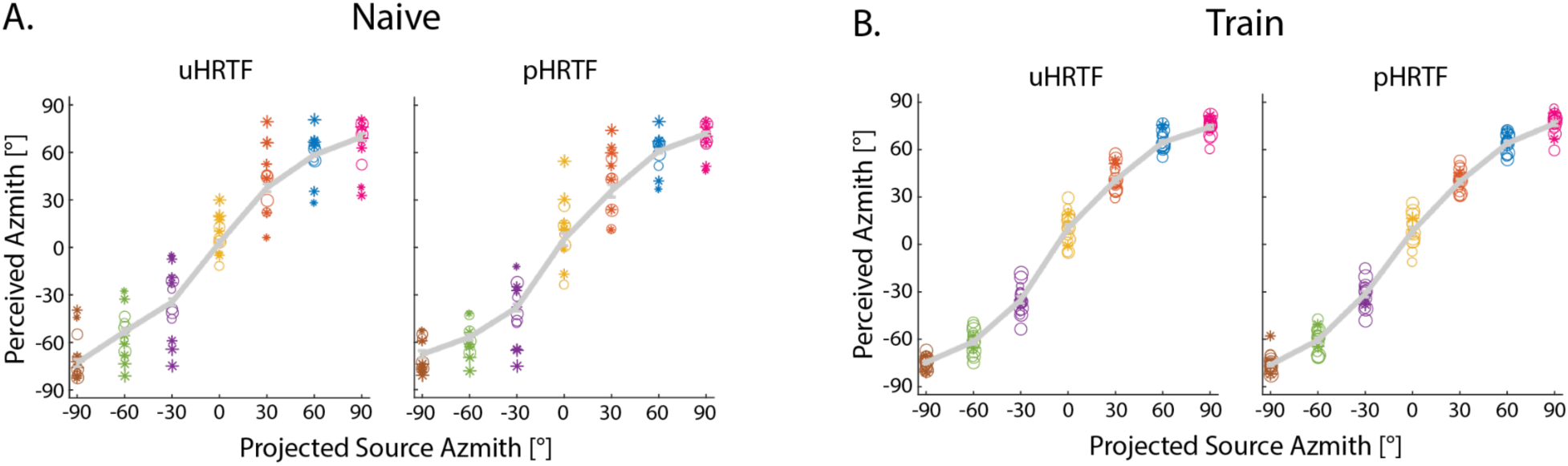
Localization of performance of azimuth as a function of projected source azimuth for (A) naive and (B) trained listeners, collapsed across all elevations. Each participant performed the localization task using a generic uHRTF or their own pHRTF. Individual participants are depicted as hollow circles (sensitive listeners) and asterisks (non-sensitive listeners), staggered from left to right and indexed by size. The solid gray lines represent the group averages for sensitive listeners. Error bars represent standard error.

The LME model for all participants showed participants were sensitive to source azimuth, as expected (α_y_*_1_*= 0.95, p < 0.001). There was a significant effect of group (α_y_*_4_* = −9.65, p < 0.001) and azimuth * group interaction (α_y_*_1_* ∗ α_y_*_4_* = −0.19, p < 0.001), suggesting an effect of training. Elevation did not significantly affect azimuth perception (α_y*2*_ = −0.04, p = 0.26). Finally, we observed no differences in which HRTF was used (α_y*3*_ = −0.82, p = 0.71). The model for only sensitive listeners showed a similar result, and the LME parameters are listed in Table S1. There was a significant effect of azimuth (α_y*1*_= 0.92, p < 0.001), as well as an azimuth * group interaction (α_y_*_1_* ∗ α_y_*_4_* = −0.16, p < 0.001). Elevation and HRTF did not significantly affect azimuth perception (α_y*2*_= −0.04, p = 0.45; α_y*3*_ = - 2.84, p = 0.41). No other model terms were significant.

### Elevation Localization

Figure 4 shows the localization of performance of elevation, collapsed across azimuths, across the two groups (“**naive”** and “**train”**) and HTRF conditions (uHRTF and pHRTF). Individual participants are depicted as hollow circles (sensitive listeners) and asterisks (non-sensitive listeners), with each participant staggered from left to right and indexed by size. The solid circles depict the group averages of the sensitive listeners.

**Figure 4.**
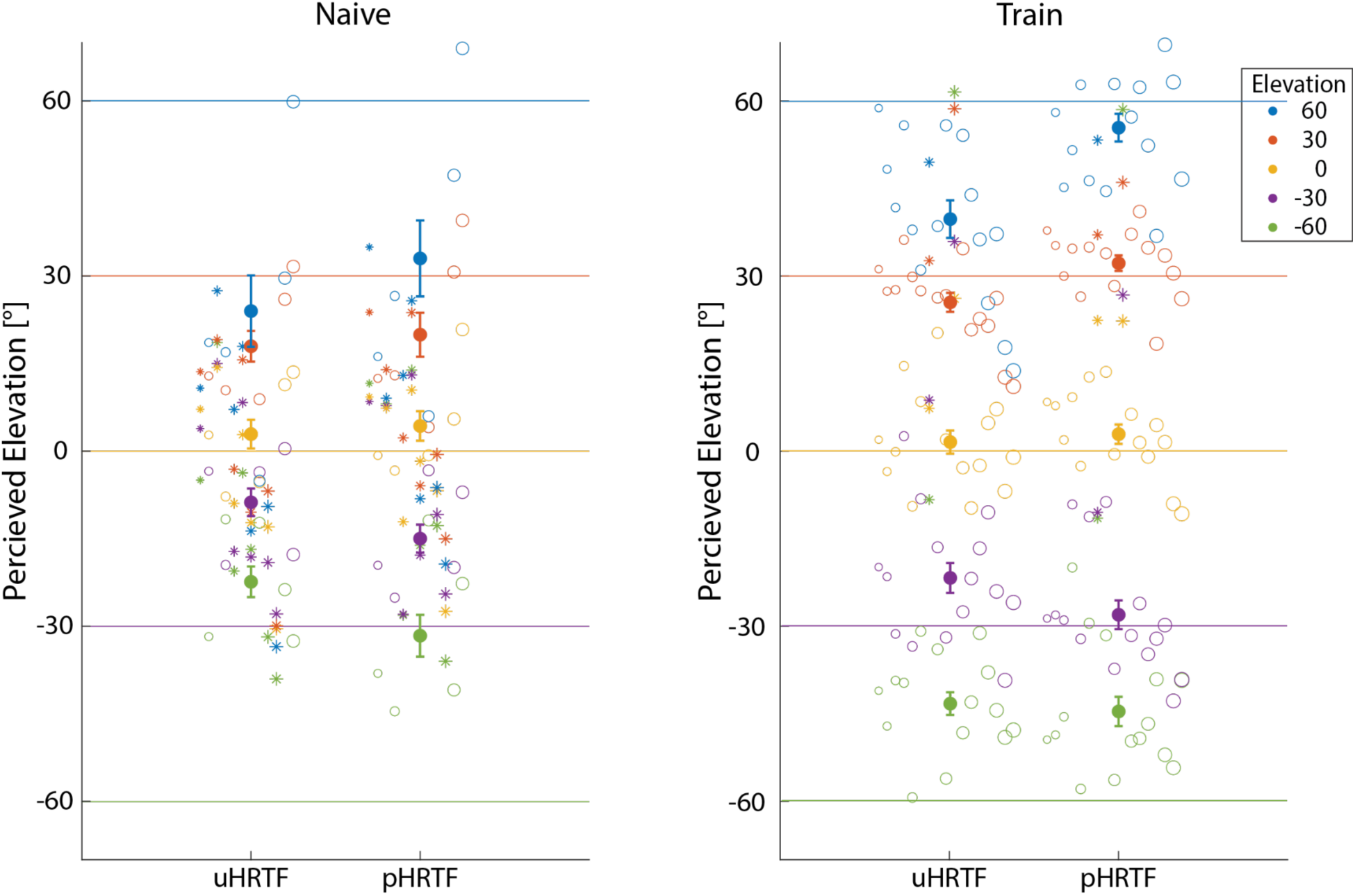
Localization of performance of elevation for (A) naive and (B) trained listeners, collapsed across all azimuths. Individual participants are depicted as hollow circles (sensitive listeners) and asterisks (non-sensitive listeners), staggered from left to right and indexed by size. The solid colored dots represent the group averages for sensitive listeners. Error bars represent standard error.

The LME model for all participants showed that participants were sensitive to source elevation (α*_y2_*= 0.25, p < 0.001). There was also a significant effect of azimuth on elevation perception (α*_y1_*= 0.02, p = 0.01). Additionally, there were significant elevation * HRTF (α*_y2_*\*α*_y3_*= 0.09, p < 0.001), elevation * group (α*_y2_*\*α*_y4_*= 0.41, p < 0.001), and elevation * HRTF * group (α*_y2_*\*α*_y3_*\*α*_y4_*= 0.05, p = 0.04) interactions, suggesting that HRTF personalization and training improved elevation perception, but more so at extreme elevation locations, and personalization benefit may increase with training. None of the other effects or interactions were significant. The model for only sensitive listeners showed a similar result, and the LME parameters are displayed in Table S2. There were significant effects of source elevation (α*_y2_*= 0.41, p < 0.001) and source azimuth (α*_y1_*= 0.03, p < 0.001). The elevation * HRTF (α*_y2_*\*α*_y3_*= 0.19, p < 0.001), elevation * group (α*_y2_*\*α*_y4_*= 0.30, p < 0.001), and HRTF * group (α*_y3_*\*α*_y4_*= 4.86, p = 0.001) interactions were also significant.

### Arc Errors

When modeling all participants, the model shows an average arc error of 49.77°, as indicated by the intercept term (β_yo_; p < 0.001). HRTF and group had significant effects, in particular, personalization decreased arc error by an average of 5.05° (α_y3_; p < 0.001), and the “**train**” group had a decreased error of 17.01° compared to the “**naive**” group (α_y*3*_; p < 0.001). Elevation significantly increased arc error, but the effect was mild (α_y*2*_ = 0.03, p = 0.02). There was also a significant elevation * group interaction (α_y*2*_*α_y*4*_= −0.07, p < 0.001) and elevation * HRTF * group interaction (α_y*2*_*α_y*3*_*α_y*4*_= −0.06, p = 0.03). No other terms were significant. In the model with sensitive listeners only, the average arc error was 38.04° (β_y*0*_; p < 0.001). HRTF personalization decreased the arc error by an average of 7.08° (α_y*3*_; p < 0.001). The “**train**” group had an average 8.02° smaller error compared to the “**naive**” group (α_y*3*_; p = 0.03). All other model terms had minimal effect on arc error. The root mean square arc errors for each HRTF condition in both groups are depicted in Figure 5A.

**Figure 5.**
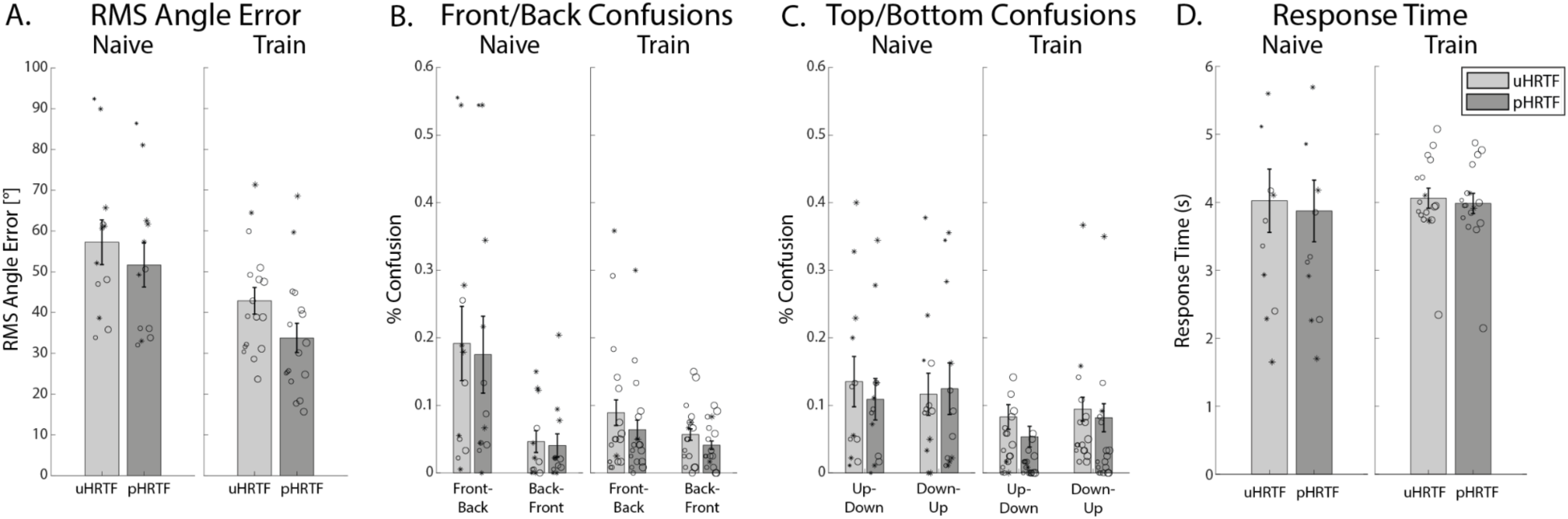
(A) Root mean squared angle error, (B) front/back confusions, (C) top/bottom confusions, (D) average response time of participants in the “naive” and “train” groups, with the light and dark gray bars representing the average results from sensitive-only participants in the uHRTF and pHRTF conditions, respectively. Individual participants are depicted as hollow circles (sensitive listeners) and asterisks (non-sensitive listeners), staggered from left to right and indexed by size. The solid colored dots represent the group averages for sensitive listeners. Error bars represent standard error.

### Front/Back Confusions

Trials where sources were in the front but perceived in the back were coded as −1; trials where sources were in the back but perceived as in the front were coded as 1; and correct front/back discriminations were coded as 0. Across all sensitive and non-sensitive listeners, front-back error rate was 12.09% (SD = 14.58%) and back-front error rate was 4.68% (SD = 4.39%). LME model of all participants showed a significant bias towards confusing sounds from the front as if they were coming from the back, as indicated by the intercept (β_y*0*_ = −0.15; p < 0.001). There was also a significant effect of group (α_y*4*_= 0.11; p = 0.04). The effects of elevation, azimuth * group, and azimuth * HRTF * group were also significant, but their estimates were all between −0.001 and 0.001. Among sensitive listeners, front-back error rate was 6.77% (SD = 5.76%) and back-front error rate was 3.97% (SD = 3.58%). The LME model of only sensitive participants showed participants were more likely to confuse front sounds (β_y*0*_ = −0.08; p = 0.002). The effect of elevation, elevation * HRTF, and azimuth * elevation * group interactions were also significant, but their estimates were all between −0.001 and 0.001. The front/back confusion probability for each HRTF condition in both groups are depicted in Figure 5B.

### Top/Bottom Confusions

Trials where sources were in the top, but perceived as in the bottom were coded as −1; trials where sources were in the bottom but perceived as in the top were coded as 1; and correct top/bottom discriminations were coded as 0. Across all listeners, top-bottom error rate was 6.84% (SD = 8.85%) and bottom-top error rate was 8.81% (SD = 9.90%). LME model of all participants showed a significant effect of HRTF (α_y*3*_= 0.03; p = 0.02).

There were also significant effects of azimuth, elevation, elevation * HRTF, and elevation * group, but their estimates were all between −0.01 and 0.01. Among sensitive listeners, the average top-bottom error rate was 4.36% (SD = 3.71%) and bottom-top error rate was 5.22% (SD = 3.83%). Using only sensitive listeners, the LME model showed a significant effect of HRTF (α_y*3*_= −0.05; p < 0.001) and HRTF * group (α_y*3*_*α_y*4*_= 0.07, p < 0.001). The effects of azimuth, elevation, azimuth * group, elevation * HRTF, and elevation * group were also statistically significantly, but with small estimates between - 0.01 and 0.01. The top/bottom confusion rates for each HRTF condition in both groups are depicted in Figure 5C.

### Response Time

The average response time for all participants was 4.02 seconds (β_y*0*_; p < 0.001). There was a significant effect of HRTF (α_y*3*_= −0.14; p <0.001), which suggests that personalization decreases response time. The effect of group (α_y*4*_= 0.03; p = 0.55), and HRTF * group interaction (α_y*3*_*α_y*4*_= 0.07; p = 0.18) on response time were both not significant. All other effects were minimal, ranging between −0.01 to 0.01 seconds.

Looking at only sensitive listeners, the average response time was 4.03 seconds (β_y*0*_; p < 0.001). There was a significant effect of HRTF (α_y*3*_= −0.32; p = 0.14) and HRTF * group interaction (α_y*3*_*α_y*4*_=0.22; p = < 0.01), reflecting HRTF personalization was stronger in the “naive” group than in the “train” group. There was no effect of group (α_y*4*_= 0.04; p = 0.93). All other estimates ranged between −0.01 to 0.01. The average response times for each HRTF condition in both groups are depicted in Figure 5D.

We wondered if localization accuracy was related to the response time, as seen in the classical speed-accuracy tradeoff phenomenon (Heitz, 2014), such that participants who spent more time on the task exhibited less error. In particular, there was one participant who on average responded before the sound stopped playing (less than 2 seconds), and was classified as a non-sensitive listener. A correlation between median response time and root mean square arc error across all participants was not significant for either the uHRTF (Pearson’s r = 0.23, p = 0.23) or pHRTF (0.21, p = 0.28) conditions.

## Discussion

Our primary interest in Experiment 1 was to assess the performance of inexperienced listeners versus those that underwent a brief training session as they localized sounds rendered with a generic HRTF compared to their own HRTF in VR. Results showed that participants were able to perceive spatialized audio in a variety of locations. However, among naive listeners without training, over half of the participants were deemed non-sensitive to spatial audio, such that they were not able to localize sounds in certain azimuths or elevations, even with their own HRTF. Statistical analysis performed on all listeners and only sensitive listeners showed similar results. We found that both HRTF personalization and belonging in the **“train”** group decreased the amount of arc error and improved elevation localization. The influence of being in the **“train”** group was much stronger than that observed from HRTF personalization. Additionally, azimuth accuracy was increased in the **“train”** group, but HRTF had no significant effect on azimuth.

Among sensitive listeners, participants were overall quite accurate in localizing within the same hemisphere front and back, as well as top and bottom. Both HRTF and group had a statistically significant effect on confusions, but their overall influence was small.

### Experiment 2

Experiment 1 demonstrated that participants who underwent a training session prior to the localization task performed better than those that were completely inexperienced. The effect of HRTF personalization was statistically significant, but small. Additionally, multiple participants were unable to accurately localize spatial audio even with their own HRTF without training. Our results suggest that training may be critical to enable naive users to acclimate to spatial audio perception in VR.

However, as Experiment 1 was a between-subject design, it is subject to limitations of the unknown individual differences among participants between groups and the inability to directly observe the effect of training over time. In Experiment 2, we recruited a completely new group of naive, inexperienced participants, and tested their accuracy in localizing sounds before and after training on two separate sessions. For comparability with the previous experiment, the procedure of Session 1 was identical to the “naive” group in Experiment 1, and Session 2 was identical to the “train” group in Experiment 1. To avoid potential confounds of how well an individual’s HRTF matched that of the uHRTF (Middlebrooks & Green, 1991; Nicol et al., 2006; Stitt et al., 2019), here, all participants were tested on their own personalized pHRTF. We hypothesized that training would improve localization accuracy, particularly elevation and arc error.

Additionally, we noticed that in Experiment 1, participants were bad at localizing extreme low elevations (i.e., −60°), even amongst sensitive listeners in the train group. We hypothesized that this may be related to the fact that participants were seated in this experiment, making certain locations difficult to reach, due to the knees or chair limiting arm motion. In Experiment 2, we had the same participants perform the localization task in sitting as well as in standing conditions, and examined whether body position had an effect on localization performance. If the lack of low elevation responses was due to body position, participants would be better at localizing lower elevations when standing.

## Methods

### Participants

Sixteen new participants were recruited for the multi-session study (ages: 26-43, mean = 32.50, SD = 6.25; 6 males, 10 females). Three additional participants were excluded due to technical difficulties, and one additional participant was excluded as they could not complete the study. All participants self-reported to have normal hearing and had their HRTF measured in our anechoic chamber within the past two years. The number of days between the first and second session ranged between 1-34 days, mean = 9.06, SD = 7.90.

### Experimental Design and Stimuli

In Experiment 2, participants who had not previously participated in spatial audio localization studies at Meta Reality Labs immediately underwent the localization test on the first session without any prior training, which was identical to the “**naive**” group in Experiment 1. In their second session, they underwent the same protocol as the “**train**” group in Experiment 1, where they learned to localize spatial audio with visual feedback for one block prior to completing the localization test.

We tested the participants in both sitting and standing conditions, to examine whether body position had an effect on the extreme low elevations (i.e., −60°). If the lack of sensitivity to low elevations was due to the knee or chair limiting arm motion, we would expect that localization for those regions would improve in the standing condition. In the first session, participants performed two blocks sitting and standing each. In the second session, participants also performed two blocks in each body position, with an additional training session before each body position condition. The body position order was counterbalanced across participants. Given the small effect of personalization observed in Experiment 1, sounds in the current experiment were rendered using only the participant’s own HRTF. This resulted in a 2 × 2 factorial design, of Session (1 vs. 2) and Body Position (sitting vs. standing).

### Statistical analysis

Statistical analysis was the same as Experiment 1. We defined sensitive listeners as those who were able to localize sounds by Session 2 with their own HRTF from all four quadrants defined in Experiment 1. The LME model is described as in Equation 2. We fitted the measurements of interest by maximum likelihood, using full interaction models, with Azimuth (α*_Y1_*), and Elevation (α*_Y2_*), Body Position (α*_Y3_*), Session (α*_Y4_*) as fixed effects. Azimuths behind the participant were folded into the front. Individual differences in the overall range of elevations were modeled via random effect β_y0_.

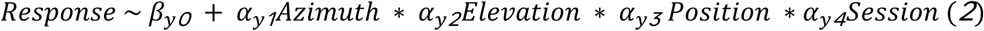

We defined sensitive listeners in an unbiased manner, using the four quadrants in space defined in Experiment 1. This classification of sensitive and non-sensitive sessions (either condition) matched that of human classification (Cohen’s Kappa = 0.95).

## Results

### Participant characteristics

In Session 1, in the sitting condition, 6 out of the 16 participants (37.5%) were labeled as non-sensitive, whereas in the standing condition, 8 out of 16 participants (50%) were non-sensitive. However, we note that in the human classification, all of these 8 participants were considered non-sensitive in both sitting and standing conditions in Session 1. In Session 2, in both sitting and standing conditions, 5 out of the 16 participants remained non-sensitive to spatial audio (31.25%). The 11 remaining participants were used in the group analysis of sensitive-only participants.

To examine whether the distribution of non-sensitive listeners before and after training in Experiment 2 were different from Experiment 1, chi-squared tests were conducted. By comparing the portion of non-sensitive participants in the **“naive”** group Experiment 1 with that of Experiment 2 Session 1, we found no significant association between listener sensitivity and experiment (χ² = 0.19, p = 0.66). Similarly, the comparison of the portion of non-sensitive participants in the **“train”** group Experiment 1 with that of Experiment 2 Session 2 showed no significant association between listener sensitivity and experiment (χ² = 1.87, p = 0.17). These results suggest that the portion of non-sensitive listeners before and after training are comparable in both experiments.

A large degree of variation was observed across participants. Among those participants who were non-sensitive to spatial audio, common errors include (1) lacked elevation sensitivity (all sounds around horizon; N=1), (2) perceived sounds mostly around or above horizon (N=5), (3) perceived a large portion of sounds from the back (N=2), and (4) discriminated sounds mostly only left versus right (N=4). Figure 6 illustrates some examples of individual participants who showed significant improvement over the course of the two sessions. OUP216 and HWN927 showed bias of localizing sounds above horizon in Session 1, and were therefore labeled non-sensitive; but in Session 2, both participants were able to resolve up-down confusions, and localize sounds both above and below horizon. ACI964 showed a tendency to localize sounds lateralized in the back in Session 1, therefore, also labeled as non-sensitive, but was able to correctly identify more sounds in the front in Session 2. DAU102 showed the most improvement, being able to distinguish only left and right locations in the horizontal plane in Session 1; was later able to perceive various azimuths and elevations after training in Session 2, although still lacked sensitivity to higher elevations.

**Figure 6.**
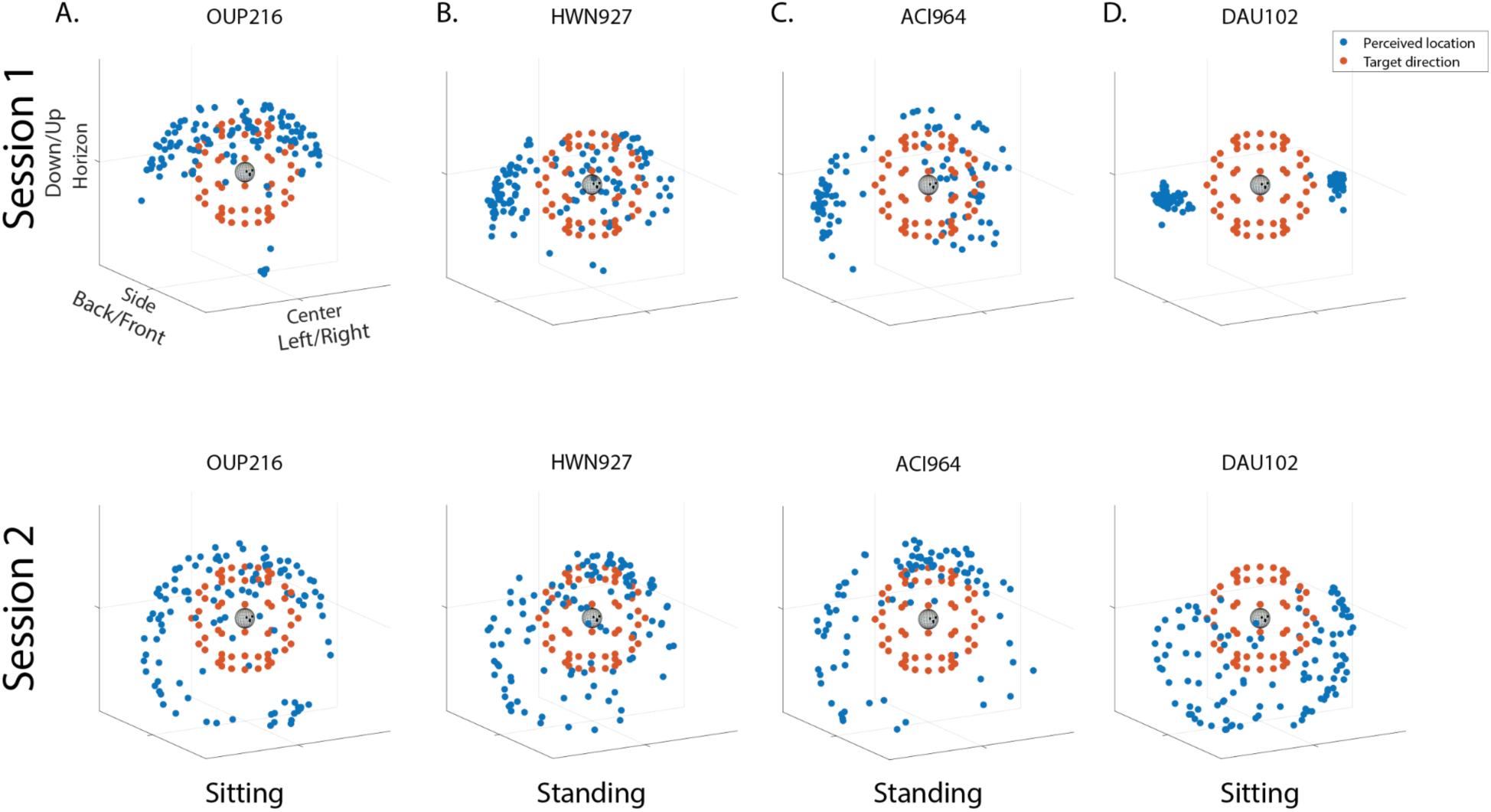
Example responses of four participants in Session 1 and Session 2. The red dots indicate the direction of the source source (scaled at 20% of the original distance for visualization). The blue dots indicate participants’ responses, with each dot representing a response for each trial. (A) Participant OUP216 showed an upward bias in their responses in Session 1, but was able to perceive sound in both top and bottom hemispaces in Session 2; (B) HWN927 lacked elevation sensitivity and perceived sounds largely around or above horizon in Session 1, but showed increased elevation sensitivity in Session 2; (C) ACI964 showed a tendency to localize sounds lateralized in the back in Session 1, but their responses were more distributed across various azimuths in Session 2; (D) DAU102 was only able to discriminate sounds as left versus right directions in Session 1 with little or no elevation perception, but was able to perceive various azimuths and elevations in Session 2, albeit still lacking sensitivity to higher elevations.

### Azimuth Localization

Figure 7 illustrates the localization of performance of azimuth, collapsed across elevations, in the sitting and standing conditions, in Session 1 and Session 2. Individual participants are depicted as hollow circles (sensitive listeners) and asterisks (non-sensitive listeners), with each participant staggered from left to right and indexed by size. Group averages of the sensitive listeners are illustrated with the solid gray line.

**Figure 7.**
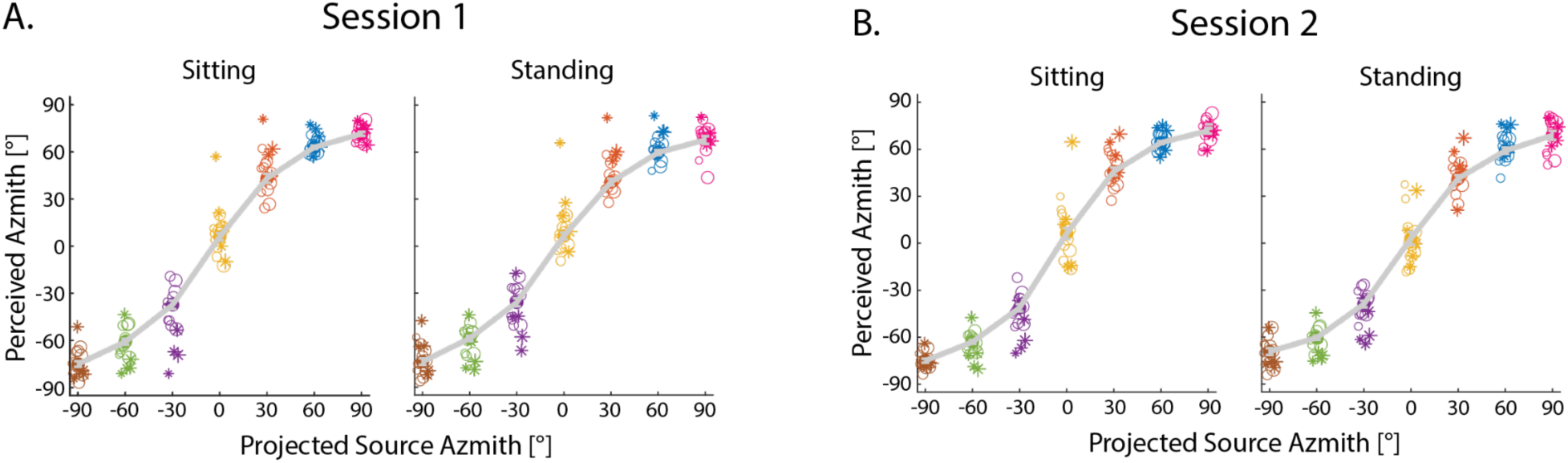
Localization of performance of azimuth as a function of projected source azimuth for (A) Session 1 and (B) Session 2 of the same participants, collapsed across all elevations, when they performed the task in sitting and standing positions. Individual participants are depicted as hollow circles (sensitive listeners) and asterisks (non-sensitive listeners), staggered from left to right and indexed by size. The solid gray lines represent the group averages for sensitive listeners. Error bars represent standard error.

The LME model for all participants showed participants were sensitive to source azimuth, as expected (α_y_*_1_*= 0.77, p < 0.001). And the model using only sensitive participants showed similar results (α_y_*_1_*= 0.75, p < 0.001). We observed an azimuth * elevation interaction in both models, but the estimates were small (α_y_*_1_*\*α_y_*_2_* < −0.001). There were no significant effects or interactions of session or body position in either model, suggesting that training and standing did not improve azimuth localization. The model coefficients for sensitive listeners are shown in Table S3.

### Elevation Localization

Figure 8 shows the localization of performance of elevation, collapsed across azimuths, across the two sessions, while performing the task sitting and standing. Individual participants are depicted as hollow circles (sensitive listeners) and asterisks (non-sensitive listeners), with each participant staggered from left to right and indexed by size. The solid circles depict the group averages of the sensitive listeners.

**Figure 8.**
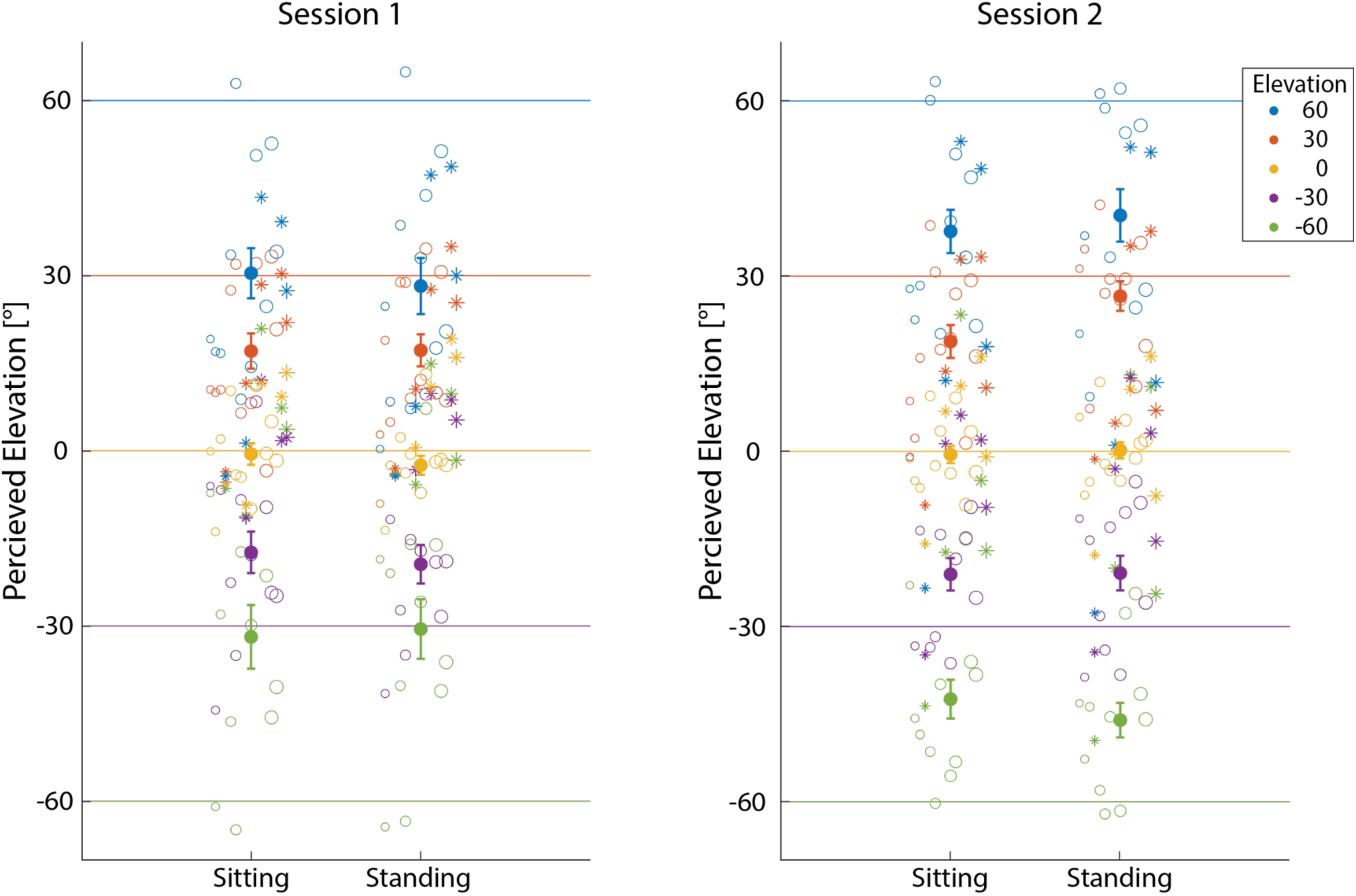
Localization of performance of elevation for (A) Session 1 and (B) Session 2, in sitting and standing positions, collapsed across all azimuths. Individual participants are depicted as hollow circles (sensitive listeners) and asterisks (non-sensitive listeners), staggered from left to right and indexed by size. The solid colored dots represent the group averages for sensitive listeners. Error bars represent standard error.

The LME model for all participants showed that elevation perception increased as a function of source elevation (α_y_*_2_*= 0.42, p < 0.001). Furthermore, we observed a significant effect of session (α_y_*_4_*= −2.51, p < 0.01), elevation * session interaction (α_y_*_2_*\*α_y_*_4_*= 0.13, p < 0.001), and elevation * position * elevation interaction (α_y_*_2_*\*α_y_*_3_*\*α_y_*_4_*= 0.06, p < 0.05). These results suggest that training increased the accuracy as well as dynamic range of elevation perception. No other terms in the model were significant.

Modeling only sensitive listeners showed a similar result. Participants were sensitive to elevation (α_y*2*_= 0.53, p < 0.001). There was a significant elevation * session interaction (α_y_*_2_*\*α_y_*_4_*= 0.14, p < 0.001) and elevation * position * session interaction (α_y_*_2_*\*α_y_*_3_*\*α_y_*_4_*= 0.10, p < 0.01), suggesting that training increased the range of more extreme elevations. The model terms are listed in Table S4.

### Arc Errors

Across all participants, the model shows an average arc error of 39.12°, as indicated by the intercept term (β_y*0*_; p < 0.001). There was a significant effect of session (α_y*4*_= −4.14, p < 0.001), which reflects that arc error decreased in Session 2. All other effects were less than 0.1°. The model using only sensitive participants, the average arc error is 34.10° (β_y*0*_; p < 0.001). Similarly, there was a significant effect of session (α_y_*_4_*= −4.24, p < 0.001) and all other effects were less than 0.1°. Figure 9A illustrates the root mean square arc errors for each session in the sitting and standing condition.

**Figure 9.**
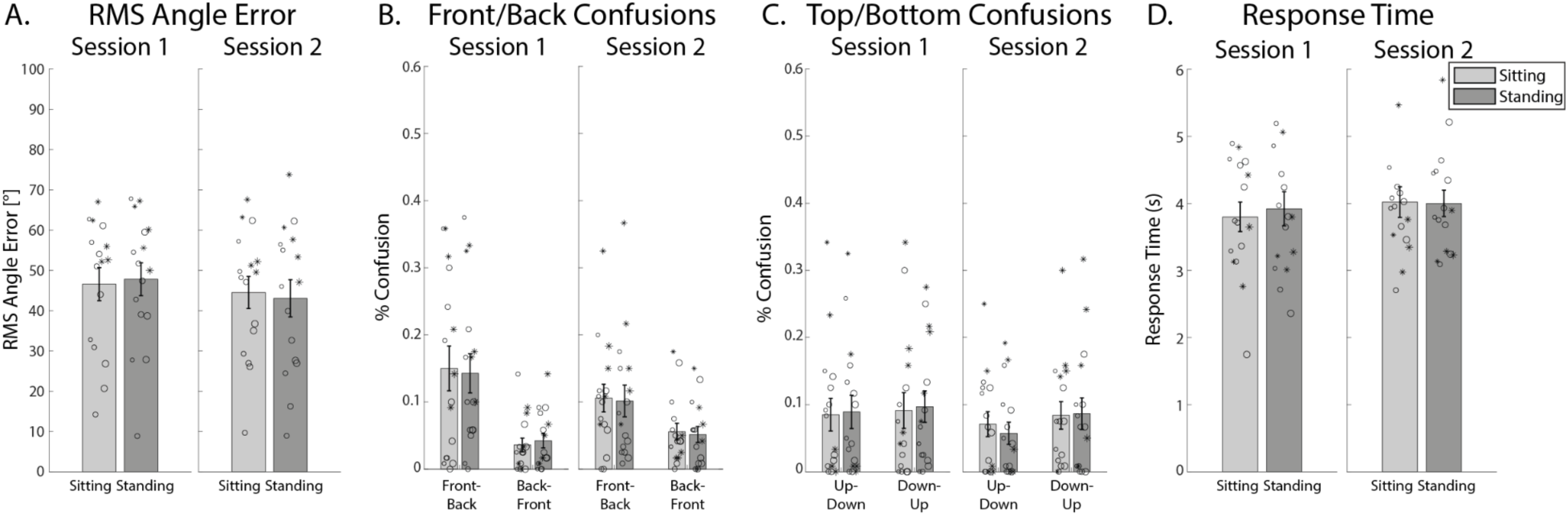
(A) Root mean squared angle error, (B) front/back confusions, (C) top/bottom confusions, (D) average response time of participants in the “naive” and “train” groups, with the light and dark gray bars representing the average results from sensitive-only participants in Session 1 and Session 2, respectively. Individual participants are depicted as hollow circles (sensitive listeners) and asterisks (non-sensitive listeners), staggered from left to right and indexed by size. The solid colored dots represent the group averages for sensitive listeners. Error bars represent standard error.

### Front/Back Confusions

Across all sensitive and non-sensitive listeners, front-back error rate was 12.50% (SD = 8.76%) and back-front error rate was 4.64% (SD = 3.21%). The LME model of all participants showed a significant bias towards confusing sounds from the front as if they were coming from the back, as indicated by the intercept (β_y_*_0_* = −0.12; p < 0.001). There was also a significant effect of session (α_y_*_4_*= 0.06, p < 0.001), reflecting a decrease in the backward bias in Session 2. There was also an effect of azimuth and elevation were also significant, but both were less than 0.001. Analyzing only sensitive listeners, the front-back error rate was 9.17% (SD = 7.06%) and back-front error rate was 4.17% (SD = 3.33%). LME modeling of the only sensitive participants also showed a significant bias towards confusing sounds from the front as if they were coming from the back (β_y*0*_ = - 0.09; p < 0.001), which decreased in Session 2 (α_y_*_4_*= 0.06, p < 0.001). The effect of azimuth was significant, but the effect was again below 0.001. The front/back confusion rates for each session and body position are shown in Figure 9B.

### Top/Bottom Confusions

Across all listeners, top-bottom error rate was 7.55% (SD = 8.03%) and bottom-top error rate was 8.96% (SD = 8.55%). The LME model of all participants showed no effect of body position (α_y_*_3_*= −0.001, p = 0.95) or session (α_y_*_4_*= 0.01, p = 0.46). All the terms that were statistically significant had small estimates, between −0.01 and 0.001. Among sensitive listeners, the average top-bottom error rate was 6.59% (SD = 5.51%) and bottom-top error rate was 5.76% (SD = 5.65%). Again, there was no significant effect of body position (α_y*3*_= −0.02, p = 0.19) or session (α_y*4*_= −0.01, p = 0.67). However, there was a body position * session interaction (α_y_*_3_*\*α_y_*_4_*= 0.04, p = 0.04), reflecting the decrease in confusions at lower elevations in Session 2. The top/bottom confusion rates for each session and body position are shown in Figure 9C.

### Response Time

The average response time for all participants was 3.79 seconds (β_y*0*_; p < 0.001). There was a main effect of azimuth (α_y*1*_< 0.001, p < 0.01), position (α_y*3*_= 0.12, p < 0.001), and session (α_y_*_4_*= 0.22, p < 0.001). There was also a position * session interaction (α_y_*_3_*\*α_y_*_4_*= - 0.14, p < 0.001). No other model terms were significant. We noted one participant was responding before the stimulus completely stopped in their Session 1 sitting condition (i.e., their average response time was less than 2 seconds). However, that participant was classified as a sensitive listener, such that their rapid response may reflect they were able to quickly identify the sound location. The result of only sensitive listeners was similar.

The average response time was 3.81 seconds (β_y*0*_; p < 0.001). There was a main effect of azimuth (α_y_*_1_*< 0.001, p = 0.01), position (α_y_*_3_*= 0.21, p < 0.001), and session (α_y_*_4_*= 0.29, p < 0.001), as well as a position * session interaction (α_y*3*_*α_y*4*_= −0.27, p < 0.001). The response times for each session and body position are shown in Figure 9D.

We explored whether there was a correlation between median response time and root mean square arc error across participants. We did not find any significant correlation between response time and error in either session or body position conditions (max Pearson’s r = 0.45, min p = 0.08).

## Discussion

Experiment 2 investigated the effect of training on the ability to localize spatial audio using participant’s own HRTF across two sessions. Results showed that training decreased overall arc errors and improved elevation localization. Training also decreased front-to-back and bottom-to-top confusions, especially among sensitive listeners. These results are consistent with that of Experiment 1, showing the benefit of training on sound localization, but within individual participants.

Across the two sessions, we were able to characterize systematic errors in localization, as well as observe improvements within subjects. In particular, three originally non-sensitive listeners, who were unable to perceive spatial audio from certain directions, became sensitive to spatial audio after training. We also observed significant improvements in participants who still remained not completely sensitive, but showed increases in the range of directions they perceived spatial audio.

Sitting and standing positions did not affect localization accuracy. This suggests that the lack of low elevation perception in Experiment 1 was not due to mobility difficulties while performing the localization task sitting in a chair. In both Experiment 1 and Experiment 2, we observed large individual differences in perceiving extreme elevations, with some participants being able to localize the +/-60° sound source accurately, others showed a compression in their responses.

Azimuth, body position, and session had an effect on reaction time. However, there was no significant correlation between response time and localization error across participants, suggesting that individual differences may not simply be a speed-accuracy tradeoff.

## General Discussion

The current study reported the performance of sound localization of naive participants and the effects of (1) training, (2) HRTF personalization, and (3) body position, across two experiments. We intended to evaluate which variables matter and to what degree when rendering realistic spatial audio in virtual reality environments. Our findings suggest that training had a significant impact on localization accuracy, whereas HRTF personalization and body position had small and no significant impact, respectively.

In Experiment 1, we examined two separate groups of participants that were either naive or underwent a brief training block with visual feedback in their ability to localize spatial audio using a generic uHRTF versus their personal pHRTF. Although there was an effect of personalization, the effect was small, and mainly affected the vertical dimension at extreme elevations. These results echo previous studies suggesting that HRTF personalization may not be necessary for users to experience accurate spatial audio, and that generic HRTFs may be adequate for many if not most use cases in VR (Berger et al., 2018; Poirier-Quinot & Katz, 2020; Rummukainen et al., 2021). This contrasts with some other studies (Jenny & Reuter, 2020; Middlebrooks, 1999; Middlebrooks et al., 2000) where the difference in performance using a generic HRTF appeared much larger. The selection of non-individualized HRTFs used in different studies may have led to different results. The degree of similarity between an individual’s HRTF and the non-individualized HRTF can also affect localization accuracy, such that a well-matched HRTF can provide good performance without training (Iida et al., 2014; Lladó, Barumerli, et al., 2024; Poirier-Quinot & Katz, 2021). Some researchers have proposed an intermediate solution to personalization, such as selecting a best-matched HRTF among a standardized set for each individual (Lladó, Pollack, et al., 2024) or predicting an individual HRTF through anthropometric features taken by a video (Brinkmann et al., 2019; Prepeliţă et al., 2020).

Another approach may be to build a good generic, i.e., “universal” HRTF that fits the majority of users. The current uHRTF within the Meta XR Audio SDK has shown improved spatial audio experience, including an 81% increase in localization accuracy, compared to the legacy model^1^.

Across Experiments 1 and 2, the most striking finding may be the effect of training. Non-sensitive listeners were characterized by their inability to perceive sound coming from at least one of four critical quadrants of space (top-left, top-right, bottom-left, or bottom-right). In Experiment 1, among the **“naive”** participants, the majority of participants were regarded as non-sensitive listeners. However, in the **“train”** group, although there remained some non-sensitive participants, most participants were able to accurately perceive spatial audio. Experiment 2 replicated the effect of training found in Experiment 1 with participants’ own HRTFs, but using a within-subjects design across two sessions. Before training, half of the naive participants showed a lack of sensitivity to spatial audio. In Session 2, after training, participants showed an overall improvement in localization accuracy, as indicated by a larger elevation range and decreased confusions. In particular, some of the non-sensitive listeners in Session 1 showed large improvements by Session 2, and were able to perceive audio in spatial regions where they originally showed absent responses. These results suggest that HRTFs alone, even personalized ones, may not enable users to immediately experience spatial audio, however, a short training session can make a significant difference as to whether listeners can experience spatial audio at all. Our results are consistent with previous findings suggesting the brain is adaptive in learning the mapping between acoustic cues and spatial locations (Barumerli et al., 2023; Carlile, 2014; King et al., 2001). Furthermore, visual and sensorimotor information from the training experience may allow participants to learn about the statistics of sound distribution and therefore adjust their responses to the range of elevations that they expect target sounds to occur (Barumerli et al., 2023; Dahmen et al., 2010; Ege et al., 2019). We note that some participants remain non-sensitive even after training in Session 2, suggesting large individual differences. While it is possible that these participants may benefit from longer training sessions (Stitt et al., 2019), it is beyond the scope of this study. Additionally, as participants did not perform the task using real-world speakers, it is unknown whether non-sensitive participants themselves were poor at localizing sounds, or whether our rendering approaches may not be sufficient to produce high fidelity audio. Nonetheless, these results highlight the importance of training in localizing sounds in VR.

Experiment 2 additionally tested whether the user was sitting or standing may affect localization accuracy, but we found very little effect of body position. This was driven by the lack of sensitivity to extreme low elevations (i.e., −60°) in the group average in Experiment 1. We hypothesized that if the lack of elevation sensitivity was due to low elevations being more physically difficult to reach in a sitting position, participants should show improved responses in the low elevation while standing. However, we found no significant differences at −60° in the sitting versus standing positions in Experiment 2, in either session. By plotting responses of individual participants, we note there is a large degree of variability across participants in their ability to localize extreme elevations, both high and low. This bias towards the center when localizing sounds in the vertical plane has been documented in previous studies (Carlile, S., Leong, P., & Hyams, S., 1997; Dobreva et al., 2011). However, the cause of this bias is currently unknown. One possibility may be the limitations in our sound rendering devices, such as the range of frequency output, digital artifact, imperfect headphone equalization,etc. Another possibility for the lack of sensitivity towards extreme elevations may be due to the mismatch between the anechoic rendering and prior expectations of floor reflections (Traer & McDermott, 2016; Wendt et al., 2017). Previous research has shown that adding room acoustics increased the perceived elevation range (Wendt et al., 2017) and accelerated HRTF learning (Poirier-Quinot & Katz, 2021). Following this conjecture, HRTFs alone may not be sufficient to reproduce a realistic spatial scene, and other factors such as room acoustics and reverberation need to be considered.

In summary, the current study extensively assessed three major factors that have been thought to influence spatial audio perception in VR. We found that HRTF personalization and body position had little effect on localization performance, however, a brief training session on visual feedback can greatly improve performance. In most VR scenarios, the spatial audio is accompanied by a visual object, such as an avatar speaking or targets in flight (Geronazzo et al., 2018; Poirier-Quinot & Katz, 2020). On one hand, acute localization will be facilitated by ventriloquism (Howard & Templeton, 1966); on the other hand, this audiovisual coupling can quickly help participants learn the spatialization of HRTFs (Berger et al., 2018). We therefore suggest that with training, generic HRTFs may be sufficient for most use cases in VR (Berger et al., 2018; Lladó, Pollack, et al., 2024; Mendonça et al., 2012).

## Acknowledgements

We would like to thank research scientists Andrew Frederick Francl, W. Owen Brimijoin, and Ishwarya Ananthabhotla for helpful discussions. We would like to thank software engineers Alex Bezugly, Eli Stine, and Christopher Poovey for making the Unity app for running this experiment. We would like to thank the data collection team: Prachetas Gumaste, Sharon Wong, Christopher Pino, José Aguayo-Barragan, Jessica Liberman, Kevin Scheumann for help with data collection. We would like to thank Andrew Luck, Phillip Robinson, and Ravish Mehra for overseeing the project.

## Supplementary Materials

**Table S1.** Results of LME model for azimuth localization for sensitive listeners.

| Term |  | Estimate | SE | t-value | p-value |  |
| --- | --- | --- | --- | --- | --- | --- |
| Intercept | $\beta_{y0}$ | -4.39 | 2.44 | -1.79 | 0.07 | |
| Azimuth | $\alpha_{y1}$ | 0.92 | 0.03 | 34.16 | 0.00 | *** |
| Elevation | $\alpha_{y2}$ | -0.04 | 0.06 | -0.75 | 0.45 | |
| HRTF | $\alpha_{y3}$ | -2.84 | 3.46 | -0.82 | 0.41 | |
| Group | $\alpha_{y4}$ | -4.12 | 2.95 | -1.40 | 0.16 | |
| Azimuth*Elevation | $\alpha_{y1}*\alpha_{y2}$ | 0.00 | 0.00 | -0.70 | 0.49 | |
| Azimuth*HRTF | $\alpha_{y1}*\alpha_{y3}$ | 0.00 | 0.04 | -0.12 | 0.91 | |
| Azimuth*Group | $\alpha_{y1}*\alpha_{y4}$ | -0.16 | 0.03 | -5.16 | 0.00 | *** |
| Elevation*HRTF | $\alpha_{y2}*\alpha_{y3}$ | -0.01 | 0.08 | -0.14 | 0.89 | |
| Elevation*Group | $\alpha_{y2}*\alpha_{y4}$ | 0.03 | 0.07 | 0.43 | 0.67 | |
| HRTF*Group | $\alpha_{y3}*\alpha_{y4}$ | 3.63 | 4.17 | 0.87 | 0.38 | |
| Azimuth*Elevation*HRTF | $\alpha_{y1}*\alpha_{y2}*\alpha_{y3}$ | 0.00 | 0.00 | -1.19 | 0.23 | |
| Azimuth*Elevation*Group | $\alpha_{y1}*\alpha_{y2}*\alpha_{y4}$ | 0.00 | 0.00 | 1.37 | 0.17 | |
| Azimuth*HRTF*Group | $\alpha_{y1}*\alpha_{y3}*\alpha_{y4}$ | -0.02 | 0.04 | -0.51 | 0.61 | |
| Elevation*HRTF*Group | $\alpha_{y2}*\alpha_{y3}*\alpha_{y4}$ | 0.01 | 0.10 | 0.05 | 0.96 | |
| Azimuth*Elevation*HRTF*Group | $\alpha_{y1}*\alpha_{y2}*\alpha_{y3}*\alpha_{y4}$ | 0.00 | 0.00 | -0.25 | 0.80 | |

Note: \*\*\* indicates $p < 0.001$ ; \*\*: $p < 0.01$ ; \*: $p < 0.05$ . A total of 5204 degrees of freedom.

**Table S2.** Results of LME model for elevation localization for sensitive listeners.

| Term |  | Estimate | SE | t-value | p-value |  |
| --- | --- | --- | --- | --- | --- | --- |
| Intercept | $\beta_{y0}$ | 3.31 | 2.81 | 1.18 | 0.24 | |
| Azimuth | $\alpha_{y1}$ | 0.03 | 0.01 | 3.63 | 0.00 | *** |
| Elevation | $\alpha_{y2}$ | 0.41 | 0.02 | 20.08 | 0.00 | *** |
| HRTF | $\alpha_{y3}$ | -1.57 | 1.23 | -1.28 | 0.20 | |
| Group | $\alpha_{y4}$ | -2.80 | 3.26 | -0.86 | 0.39 | |
| Azimuth*Elevation | $\alpha_{y1}*\alpha_{y2}$ | 0.00 | 0.00 | -0.83 | 0.41 | |
| Azimuth*HRTF | $\alpha_{y1}*\alpha_{y3}$ | -0.02 | 0.01 | -1.53 | 0.13 | |
| Azimuth*Group | $\alpha_{y1}*\alpha_{y4}$ | -0.02 | 0.01 | -2.13 | 0.03 | * |
| Elevation*HRTF | $\alpha_{y2}*\alpha_{y3}$ | 0.19 | 0.03 | 6.63 | 0.00 | *** |
| Elevation*Group | $\alpha_{y2}*\alpha_{y4}$ | 0.30 | 0.02 | 12.28 | 0.00 | *** |
| HRTF*Group | $\alpha_{y3}*\alpha_{y4}$ | 4.86 | 1.48 | 3.28 | 0.00 | *** |
| Azimuth*Elevation*HRTF | $\alpha_{y1}*\alpha_{y2}*\alpha_{y3}$ | 0.00 | 0.00 | -0.11 | 0.91 | |
| Azimuth*Elevation*Group | $\alpha_{y1}*\alpha_{y2}*\alpha_{y4}$ | 0.00 | 0.00 | -0.10 | 0.92 | |
| Azimuth*HRTF*Group | $\alpha_{y1}*\alpha_{y3}*\alpha_{y4}$ | 0.02 | 0.02 | 1.24 | 0.22 | |
| Elevation*HRTF*Group | $\alpha_{y2}*\alpha_{y3}*\alpha_{y4}$ | -0.03 | 0.03 | -0.90 | 0.37 | |
| Azimuth*Elevation*HRTF*Group | $\alpha_{y1}*\alpha_{y2}*\alpha_{y3}*\alpha_{y4}$ | 0.00 | 0.00 | -0.32 | 0.75 | |

Note: \*\*\* indicates $p < 0.001$ ; \*\*: $p < 0.01$ ; \*: $p < 0.05$ . A total of 5204 degrees of freedom.

**Table S3.** Results of LME model for azimuth localization for sensitive listeners.

| Term |  | Estimate | SE | t-value | p-value |  |
| --- | --- | --- | --- | --- | --- | --- |
| Intercept | $\beta_{y0}$ | -4.32 | 2.06 | -2.09 | 0.04 | |
| Azimuth | $\alpha_{y1}$ | 0.75 | 0.02 | 37.81 | 0.00 | *** |
| Elevation | $\alpha_{y2}$ | 0.04 | 0.05 | 0.81 | 0.42 | |
| Position | $\alpha_{y3}$ | 0.11 | 2.92 | 0.04 | 0.97 | |
| Session | $\alpha_{y4}$ | -1.63 | 2.92 | -0.56 | 0.58 | |
| Azimuth*Elevation | $\alpha_{y1}*\alpha_{y2}$ | 0.00 | 0.00 | -3.03 | 0.00 | *** |
| Azimuth*Position | $\alpha_{y1}*\alpha_{y3}$ | 0.02 | 0.03 | 0.65 | 0.51 | |
| Azimuth*Session | $\alpha_{y1}*\alpha_{y4}$ | 0.01 | 0.03 | 0.41 | 0.68 | |
| Elevation*Position | $\alpha_{y2}*\alpha_{y3}$ | 0.01 | 0.07 | 0.16 | 0.87 | |
| Elevation*Session | $\alpha_{y2}*\alpha_{y4}$ | -0.09 | 0.07 | -1.34 | 0.18 | |
| Position*Session | $\alpha_{y3}*\alpha_{y4}$ | -2.93 | 4.13 | -0.71 | 0.48 | |
| Azimuth*Elevation*Position | $\alpha_{y1}*\alpha_{y2}*\alpha_{y3}$ | 0.00 | 0.00 | 0.87 | 0.39 | |
| Azimuth*Elevation*Session | $\alpha_{y1}*\alpha_{y2}*\alpha_{y4}$ | 0.00 | 0.00 | -0.16 | 0.87 | |
| Azimuth*Position*Session | $\alpha_{y1}*\alpha_{y3}*\alpha_{y4}$ | 0.00 | 0.04 | 0.10 | 0.92 | |
| Elevation*Position*Session | $\alpha_{y2}*\alpha_{y3}*\alpha_{y4}$ | 0.01 | 0.10 | 0.14 | 0.89 | |
| Azimuth*Elevation*Position*Session | $\alpha_{y1}*\alpha_{y2}*\alpha_{y3}*\alpha_{y4}$ | 0.00 | 0.00 | 0.40 | 0.69 | |

Note: \*\*\* indicates $p < 0.001$ ; \*\*: $p < 0.01$ ; \*: $p < 0.05$ . A total of 5264 degrees of freedom.

**Table S4.** Results of LME model for elevation localization for sensitive listeners.

| Term |  | Estimate | SE | t-value | p-value |  |
| --- | --- | --- | --- | --- | --- | --- |
| Intercept | $\beta_{y0}$ | -0.02 | 2.09 | -0.01 | 0.99 | |
| Azimuth | $\alpha_{y1}$ | -0.01 | 0.01 | -1.34 | 0.18 | |
| Elevation | $\alpha_{y2}$ | 0.53 | 0.02 | 30.90 | 0.00 | *** |
| Position | $\alpha_{y3}$ | -1.01 | 1.04 | -0.97 | 0.33 | |
| Session | $\alpha_{y4}$ | -1.29 | 1.04 | -1.24 | 0.22 | |
| Azimuth*Elevation | $\alpha_{y1}*\alpha_{y2}$ | 0.00 | 0.00 | -0.80 | 0.42 | |
| Azimuth*Position | $\alpha_{y1}*\alpha_{y3}$ | 0.00 | 0.01 | 0.27 | 0.78 | |
| Azimuth*Session | $\alpha_{y1}*\alpha_{y4}$ | 0.01 | 0.01 | 0.57 | 0.57 | |
| Elevation*Position | $\alpha_{y2}*\alpha_{y3}$ | -0.02 | 0.02 | -0.90 | 0.37 | |
| Elevation*Session | $\alpha_{y2}*\alpha_{y4}$ | 0.14 | 0.02 | 5.52 | 0.00 | *** |
| Position*Session | $\alpha_{y3}*\alpha_{y4}$ | 2.62 | 1.47 | 1.79 | 0.07 | |
| Azimuth*Elevation*Position | $\alpha_{y1}*\alpha_{y2}*\alpha_{y3}$ | 0.00 | 0.00 | -0.63 | 0.53 | |
| Azimuth*Elevation*Session | $\alpha_{y1}*\alpha_{y2}*\alpha_{y4}$ | 0.00 | 0.00 | -1.93 | 0.05 | |
| Azimuth*Position*Session | $\alpha_{y1}*\alpha_{y3}*\alpha_{y4}$ | -0.01 | 0.01 | -0.51 | 0.61 | |
| Elevation*Position*Session | $\alpha_{y2}*\alpha_{y3}*\alpha_{y4}$ | 0.10 | 0.03 | 2.89 | 0.00 | *** |
| Azimuth*Elevation*Position*Session | $\alpha_{y1}*\alpha_{y2}*\alpha_{y3}*\alpha_{y4}$ | 0.00 | 0.00 | 0.73 | 0.47 | |

*Note: \*\*\* indicates $p < 0.001$ ; \*\*: $p < 0.01$ ; \*: $p < 0.05$ . A total of 5264 degrees of freedom.*

## Footnotes

1 https://developers.meta.com/horizon/blog/improve-spatial-audio-universal-hrtf-meta-quest/

